# Parameterization of cell-free systems with time-series data using KETCHUP

**DOI:** 10.1101/2025.06.06.658289

**Authors:** Mengqi Hu, S. Bilal Jilani, Daniel G. Olson, Costas D. Maranas

## Abstract

Kinetic models mechanistically link enzyme levels, metabolite concentrations, and allosteric regulation to metabolic reaction fluxes. This coupling allows for the quantitative elucidation of the dynamics of the evolution of metabolite concentrations and metabolic fluxes as a function of time. So far, most large-scale kinetic model parameterizations are carried out using mostly steady-state flux measurements supplemented with metabolomics and/or proteomics data when available. Even though the parameterized kinetic model can trace a temporal evolution of the system, lack of anchoring to temporal data reduces confidence in the dynamics predictions. Notably, the simulation of enzymatic cascade reactions requires the full description of the dynamics of the system as a steady-state is not applicable given that all measured metabolite concentrations vary with time. Here we describe how kinetic parameters fitted to the dynamics of single-enzyme assays remain accurate for the simulation of multi-enzyme cell-free systems. Herein, we demonstrate two extensions for the Kinetic Estimation Tool Capturing Heterogeneous datasets Using Pyomo (KETCHUP) software tool for parameterizing a kinetic model of the cell-free kinetics of formate dehydrogenase (FDH) and 2,3-butanediol dehydrogenase (BDH) through the use of time-course data across various initial conditions. An implemented extension of KETCHUP allowing for the reconciliation of measurement time-lag errors present in datasets was used to parameterize multiple datasets. By combining the kinetic parameters identified by the FDH and BDH assays, accurate simulation of the binary FDH-BDH system was achieved. KETCHUP can be accessed at https://github.com/maranasgroup/KETCHUP.

**Author Summary:** Metabolic engineering of microorganisms offers a sustainable and renewable alternative for industrial scale production of commodity chemicals. Large-scale metabolic stoichiometric models enable means to elucidate an organism’s physiological behavior towards both environment and genetic perturbations and predict optimal interventions to maximize production. Kinetic models extend stoichiometric models’ capabilities in characterizing metabolic phenotypes by mathematically linking enzyme catalyzed reactions in metabolic networks as functions of metabolite concentrations, enzyme levels, and allosteric regulations. Although these kinetic descriptions improve predictive accuracy and strain design, the lack of enzyme information, raw experimental data, and high computational cost preclude the construction and parameterization of detailed large-scale kinetic models. Here, we introduce an extension of KETCHUP, an open-source semi-automatic kinetic construction and parameterization tool that utilizes steady-state data as a pilot for the parameterization of single-enzyme cell-free time-course data. We show that this tool offers comparable performance to existing tools during parameterization of single enzymes and successful recapitulation of metabolite profiles for a two-enzyme system when simulating a kinetic model with parameters from the two single-enzymes. This study showcases a high throughput pipeline that can be integrated into our framework for larger scaled networks.

## Introduction

Advances in metabolic engineering has led to many successful proof-of-concept designs and experiments for the renewable production of numerous chemicals and commodities such as biofuels [1–3], pharmaceuticals [4,5], isoprenoids [6–8], fatty acids [9,10], and organic acids [11,12]. However, the commercialization of these strain designs is hindered by low titer and yields which requires pathway optimization for scalable production [13]. These issues arise from factors such as *(i)* inherent complexities of metabolic and regulatory networks which hinder optimal and simple strain designs, *(ii)* non-native biosynthetic pathways that affect cell viability and homeostasis [14,15], and *(iii)* lack of detailed kinetic information for an organism’s key enzymes. While a “whole cell” strain is often the end goal for most metabolic engineering projects, cell-free systems (CFS) can contribute to overcoming bottlenecks during the strain and pathway design process. CFSs are fundamentally different than “whole cells” as the homeostasis requirement of living cells and most forms of protein-level feedback can be circumvented allowing for proper diagnosis of kinetic information [15]. The lack of compartmentalization results in a dilute and well-mixed reaction environment allowing for high resolution in the observation of reaction kinetics [16]. This ease of experimental design can facilitate the discovery of regulatory interactions and elucidation of resource allocations [17]. The lack of enzyme self-regeneration though requires estimation of enzyme stability to reliably elucidate catalytic rates [18].

CFS can be divided into two types based on the method of preparation: (*i*) extract-based or lysate systems, and (*ii*) purified enzyme-based systems [19]. The extract-based CFS was first used for bio-ethanol production from fermenting sugar using cell extracts from yeast [20] and later used to study the dynamics in protein synthesis where researchers engineer (*i*.*e*., DNA sequences for expression) and assembled components required for protein synthesis reactions (*i*.*e*., amino acids, energy buffers and cofactors) into a cellular lysate (*i*.*e*., cytoplasmic lysate from living cell that only contains transcription and translation machinery for protein synthesis). This cell-free metabolic engineering platform has demonstrated comparable performance with cell-based systems [21], especially in the synthesis of biologic therapeutics (*e*.*g*., antibodies [22], antimicrobial peptides [23], or vaccines [24]). The key advantage of extract-based cell-free systems is that they are unconstrained by homeostatic considerations, allowing for continuous probing over a specified time horizon. Alternatively, enzyme-based CFSs use purified enzymes for single reaction or pathway (*i*.*e*., cascade reaction) exploratory purposes. This set-up allows for flexible engineering and complete control of the reaction parameters (*i*.*e*., user-defined enzyme and metabolite concentrations) over extract-based systems which has competing reactions and toxic cofactors that hinder the targeted metabolic reaction [25,26]. For example, a system of 27 purified enzymes has been shown to convert glucose to terpenes [27] or a system of three enzymes was used to produce UDP-N-acetylglucosamine from its direct precursor N-acetylglucosamine [28]. These applications demonstrate that both extract-based and enzyme-based CFSs can be valuable tools for the exploration of alternative pathway designs. However, the number of experimental variables (*i*.*e*., initial concentrations of enzymes, substrates, and cofactors) can make it difficult to optimize performance empirically, and in some cases, small changes in initial conditions can prematurely halt the reaction. For example, enzyme cascades that use the glycolysis pathway are highly sensitive to the initial hexokinase concentration [27]. Thus, there is a need to develop predictive computational models to assist in optimizing system performance, and to test hypothesis for system malfunctions.

Multiple computational methods and models have been used for the elucidation of metabolism. The earliest methods are constraint-based methods such as flux balance analysis [29] where an organism’s phenotype is simulated based on cellular objectives (*i*.*e*., growth). Unfortunately, stoichiometric models cannot explicitly relate enzyme information or regulatory processes (*e*.*g*., inhibition), limiting the advantages afforded by CFSs (*i*.*e*., qualitative dynamic data that provides vital insight towards enzyme metabolism). Kinetic metabolic models address these shortcomings by incorporating enzyme kinetics, allosteric regulatory interactions, and enzyme and metabolite levels into reaction fluxes by offering a more comprehensive description of cell metabolism than stoichiometric models alone. These mathematical models offer a more comprehensive description of cell metabolism [30,31] than stoichiometric models and improves predictive capabilities [32–35], strain designs [36–38], and elucidation of regulatory interactions [39–41]. Several frameworks have been developed to facilitate the parametrization of large-scale kinetic models from a top-down (*e*.*g*., elementary reaction steps [42]) approach by using generalized rate laws for reactions due to lacking enzyme mechanism information. However, this generalized approach obscure details on individual enzyme mechanisms (*e*.*g*., substrate/product binding/release order), especially for allosteric regulations. For example, glucose 6-phosphate dehydrogenase binds substrates sequentially [43] whereas transketolases follow a ping-pong mechanism [44]. Despite the lack of specific enzyme information, these approaches of parameterizing kinetic models with *in vivo* datasets have successfully provided sufficiently quantitative description of cellular processes [45].

Contrary to large-scale kinetic models that commonly use a “top-down” approach for parameterization by modeling “whole cells” with steady-state *in vivo* kinetic data, small-scale kinetic models that capture transient metabolism in higher resolution require a “bottom-up” approach. These small-scale models typically are used to study cell-free systems containing either a singular enzyme or cascading enzymes that constitute the pathway of interest [46,47]. Cell-free systems provide a straightforward environment, devoid of interactions from the rest of the cellular components, for the characterization of specific enzyme activities and mechanisms.

This trait allows for the construction and validation of various mechanistic models [48]. Moreover, combining advances in rapid prototyping of individual enzymes [49,50] in CFSs and efficient parameterization algorithms help speed-up the construction of high quality large-scale kinetic models.

However, one notable consideration when characterizing enzymes in CFSs is the discrepancies often observed between *in vivo* and *in vitro* kinetic parameters, especially for turnover numbers (*i*.*e*., both k_app_ and k_cat_ values for both prokaryotes [51] and eukaryotes [52]). Fortunately, studies on CFSs that mimic *in vivo* environments [53,54] offer the means to bridge the gap between *in viv*o and *in vitro* kinetic parameter values [55]. This alignment could help improve resolution of large-scale kinetic models compared to ones that are constructed with kinetic parameters from varying environmental conditions [56]. Parameterization of large-scale kinetic models constructed with varying mechanistic rate laws require tools that allow for customized inputs. Existing kinetic parameter estimation tools such as COPASI [57], Tellurium [58], or SkiMpy [59] allow for customized kinetic rate laws but require direct integration of a systems of ordinary differential equations, which hamper model network scalability [60,61]. Scalability and rate law flexibility motivated the development of the Kinetic Estimation Tool Capturing Heterogeneous datasets Using Pyomo (KETCHUP) which semi-automatically constructs and parameterizes kinetic models [62]. The original implementation of KETCHUP was benchmarked against previous kinetic models parameterized with steady-state data for groups of genetically perturbed strains; it demonstrated improved parameterization times and quality of fit compared to their respective previous parameterization tool. In this work, we extend KETCHUP’s parameterization pipeline to the use of time-course data. Specifically, kinetic models are parameterized with *in vitro* time-course data of formate dehydrogenase (FDH) from *Candida boidinii* and 2,3-butanediol dehydrogenase (BDH) from *Serretia marcescens* using simplified Michaelis-Menten kinetics. A KETCHUP extension is used to identify systematic experimental errors arising from time-delayed measurements and propose lag-times needed to be imposed on the data to enable correct recapitulation of the fitted data. By integrating separately parameterized FDH and BDH kinetic parameter values, the experimental measured metabolite profiles for the two enzyme cell-free system FDH-BDH are recapitulated.

## Results

Three distinct time-course dataset series generated in this study (*i*.*e*., two formate dehydrogenase, FDH and one 2,3-butanediol dehydrogenase, BDH) were curated and used for the parameterization of two distinct kinetic models. Both models were constructed following their reaction mechanism depicted in Figure 1. To first assess KETCHUP capability of fitting time-course data, we parameterized a FDH model with published data (dataset series A1) and compare kinetic parameters found with their respective reported values [63]. Similar parameter fits were achieved compared to the published kinetic parameters (See Figure 2). Next, we proceeded with the parameterization of datasets generated in this study for the FDH (dataset series B1 and B2) and BDH (dataset series Z1) systems with and without time-lag adjustments. These adjustments were made to help reconcile reaction start time and reported measurement times. We observed that time-lag adjustments only improve fitting of the datasets by 15%. Finally, we examine KETCHUP’s simulation capability of using single-enzyme identified kinetic parameters to simulate a binary system in a fed-batch system. Results indicate that single-enzyme fitted kinetic parameters can be combined and used in larger kinetic models for prediction of metabolite profiles (see Figure 4).

**Figure 1.**
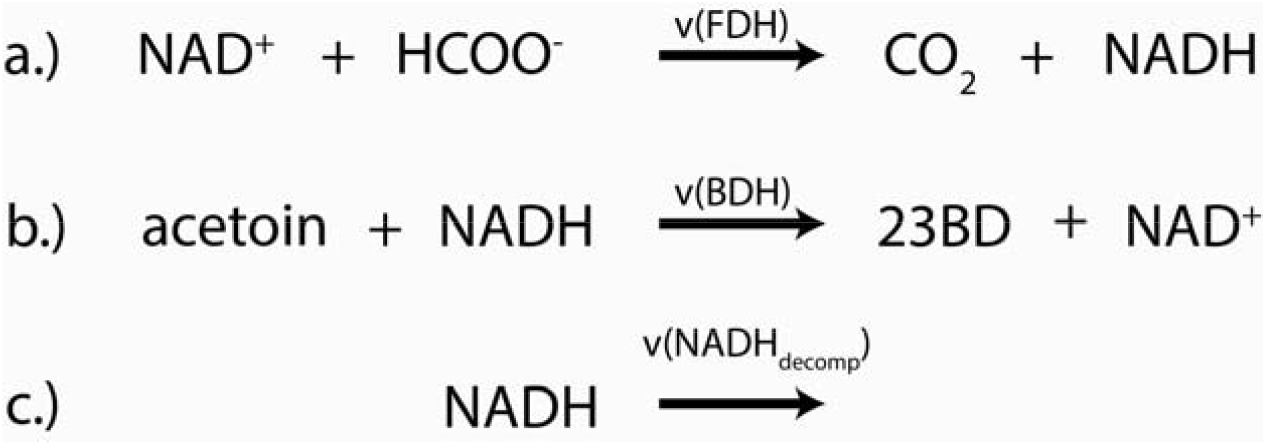
Reaction mechanisms for a) formate dehydrogenase (FDH), b) 2,3-butanediol dehydrogenase (BDH), and c) NADH decomposition reactions. Hydrogen is excluded in the kinetic model and reflected in this schematic for BDH. Rates (v) for each reaction are defined in section “model construction”

**Figure 2.**
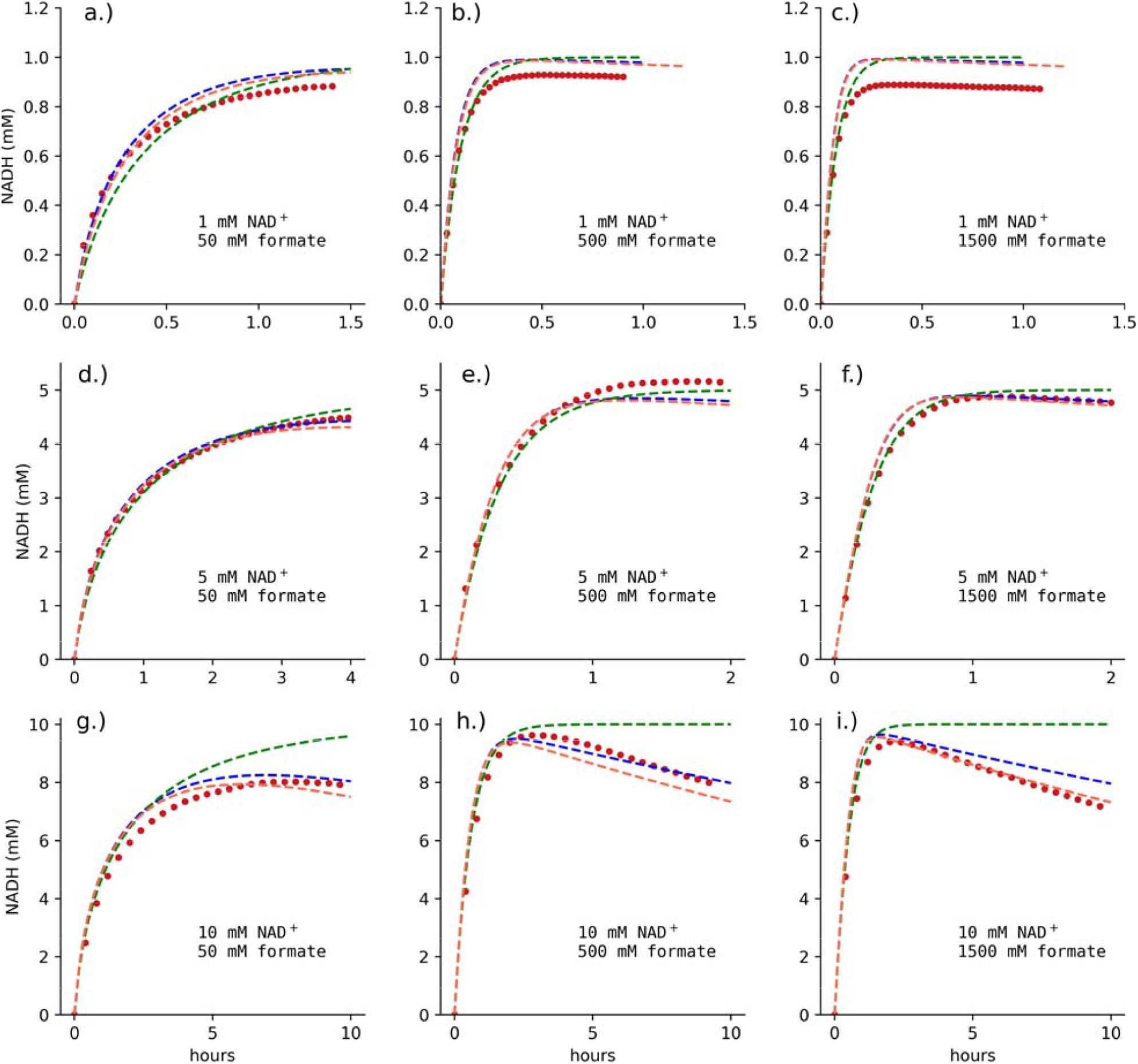
Prediction of NADH concentration over time between experimental data used in parameterization (red dots), simulation using KETCHUP-determined parameters with (dashed blue line) and without (dashed green line) NADH decomposition reaction, and previously reported fitted kinetic parameters (dashed orange line).

### Raw data filtering for parameterization

FDH time-course data with varying substrate, product, and enzyme concentrations were collected from a previous study [63] including (Dataset series A1) and two separate experiments (Datasets series B1 and B2). Due to unavailability of tabulated data from the literature study, Dataset series A1’s data points were extracted from the plot figure provided in the study using WebPlotDigitizer (automeris.io). BDH time course data with varying substrate, product, and enzyme concentrations were collected via experiments (Dataset series Z1). Experimental set-up and measurements are described in methods.

The collected raw time-course NADH concentration data were filtered by calculating a moving average of 10 datapoints per time point to smoothen out measurement noise. The data is then systematically thinned by using the Ramer-Douglas-Peucker (RDP) algorithm [64] to reduce the number of datapoints in a curve and consequently alleviate computational burden during parameterization with minimal loss of information. The threshold for the RDP algorithm for each dataset in each series is iteratively determined such that the resulting filtering would provide between 95 to 105 datapoints per dataset. This helps to normalize the error calculation for the objective function during fitting (see methods). Each raw data and corresponding filtered data were plotted (See Supplementary Figures SF1-SF3) with selected datasets being removed from further analysis if too noisy or exhibiting obviously erroneous trends (see Supplementary Materials SM2 for exclusion criteria) following consultation with the data generation team. In summary, 13, 29, and 34 datasets out of 32, 88, and 134 were filtered from Dataset series B1, B2, and Z1, respectively. The initial RDP-filtered dataset series are in Supplementary Materials SM2 while resulting curated dataset series used for parameterization can be found at https://github.com/maranasgroup/KETCHUP/tree/main/KETCHUP_time-series.

### Model construction

The FDH model was parameterized using a published simplified Michaelis-Menten style mechanistic rate law [63] (see Equation 2) where v is the reaction rate, V_max_ is maximal reaction rate, and A, B, and Q represent metabolite NAD^+^, formate, and NADH, respectively. Parameters K_M_ and K_I_ denote Michaelis-Menten and inhibitor constants, respectively. To parameterize varying enzyme concentrations V_max_ is set to be equal to a turnover number k_cat_ times the enzyme concentration E^FDH^. The BDH model was parameterized using a convenience kinetic mechanistic rate law [65] (see Equation 3), where 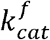 and 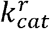 is the forward and reverse turnover number, respectively, S, P, and E^BDH^ representing metabolite acetoin and 2,3-butanediol and enzyme 2,3-butanediol dehydrogenase, respectively.

*Equation 2: Mechanistic kinetic rate law describing FDH presented by [63]*.

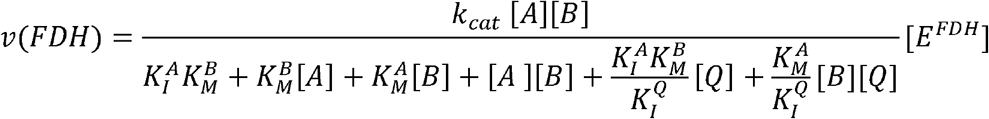

*Equation 3: BDH rate law represented in convenience kinetics presented by [65]*.

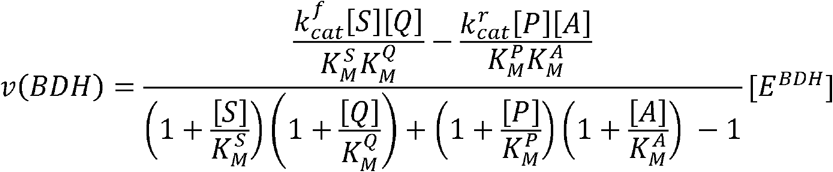

Both kinetic models include an irreversible non-enzymatic decomposition reaction [66] for cofactor NADH for a more accurate description of the reaction dynamics over a long period of time (See Equation 4) where k_dQ_ is the NADH decomposition rate constant.

*Equation 4: NADH decomposition is assumed to follow first-order kinetics as presented in [63]*.

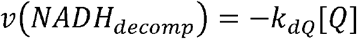

### Comparison of kinetic parameters against previous study

To benchmark KETCHUP’s extension towards parameterizing time-course data, we fitted the FDH kinetic model, containing reaction rate law for FDH and an irreversible first order NADH decomposition, with nine published datasets with varying initial NADH and formate concentrations [63]. Due to lack of tabulated datasets, the dataset series A1 was digitized to contain between 24 to 37 evenly spaced datapoints for each set. To sample across the solution space, 100 randomly initialized multi-starts were performed. All solutions reached convergence with an average solve time of 3.82 seconds. The best solution yielded an sum of squared residuals (SSR) value of 1.19 mM^2^ (similar to the SSR found while using published kinetic parameters with SSR of 1.49 mM^2^). The kinetic parameters for both KETCHUP-parameterized and published models are listed in Table 1. Except for inhibitor constant for NAD^+^, all kinetic parameters found with KETCHUP are within the same order of magnitude as published values. A slight systematic overprediction of NADH for simulations with initial concentration of 1 mM NAD^+^ results from the objective function prioritizing minimizing the error deviation between predicted and experimental values for higher NADH concentrations. This prioritization consequently results in overprediction for the low NADH concentrations. It is important to also note that the inclusion of NADH degradation reaction better explains the datasets measured over longer periods of time [63] (See Figure 2g to 2i) as the SSR increases to 3.73 mM^2^ if NADH degradation is unaccounted for in the simulation.

**Table 1.** Average kinetic parameters found during parameterization of FDH and their published.

| Kinetic parameter | Fitted values | Published values<sup>a</sup> |
| --- | --- | --- |
| $k_{dQ}$ ($\text{h}^{-1}$) | $2.39 \times 10^{-2} \pm 4.40 \times 10^{-6}$ | $3.45 \times 10^{-2}$ |
| $K_I^A$ (mM) | $5.74 \times 10^2 \pm 2.76 \times 10^2$ | $78.14 \pm 0.08$ |
| $K_i^Q$ (mM) | $7.71 \times 10^{-1} \pm 2.31 \times 10^{-2}$ | $1.18 \times 10^{-1} \pm 4.0 \times 10^{-5}$ |
| $K_M^B$ (mM) | $7.79 \times 10^{-1} \pm 1.01$ | $4.72 \times 10^{-1} \pm 7 \times 10^{-3}$ |
| $K_M^A$ (mM) | $4.02 \times 10^{-1} \pm 2.63 \times 10^{-2}$ | $3.84 \times 10^{-2} \pm 9 \times 10^{-5}$ |
| $k_{\text{cat}}$ ($\text{s}^{-1}$) | $2.36 \times 10^{-1} \pm 5.41 \times 10^{-3}$ | 0.18 |

<sup>a</sup>Schmidt et al. 2010

KETCHUP’s parameterization computation time performance for this dataset series is comparable to MATLAB’s [67] lsqnonlin function coupled with ode45 and has slightly better SSR (1.49 mM^2^ vs 1.81 mM^2^, respectively). These metrics show that despite KETCHUP’s original development for the parameterization of large-scale models using steady-state dataset, KETCHUP performs equally well to existing tools for time-course datasets (note that reported values in [63] were found using gPROMS (Version 3.0.2, Process System Enterprise Ltd., London, UK)).

### Fitting experimental datasets and time-lag considerations

Two FDH and one BDH kinetic models were parameterized with the generated herein dataset series B1, B2, and Z1, respectively. All kinetic models were first parameterized with their corresponding NADH decomposition standards to determine a range of values for *k*_*dQ*_ per dataset series before parameterization of other parameters (*i*.*e*., [1.50,3.50]×10^−4^ min^-1^, [1.11,1.77]×10^−4^ min^-1^, [1.286,1.287]×10^−3^ min^-1^ for dataset series B1, B2, and Z1, respectively). Each kinetic model was parameterized while constraining *k*_*dQ*_ to their predetermined range with 500 randomly initialized multi-starts. Performance metrics are listed in Table 2. Note that *k*_*dQ*_ for FDH experiments is one order of magnitude less than that for BDH experiments resulting from different experimental set-up (*i*.*e*., assay buffer).

**Table 2.**
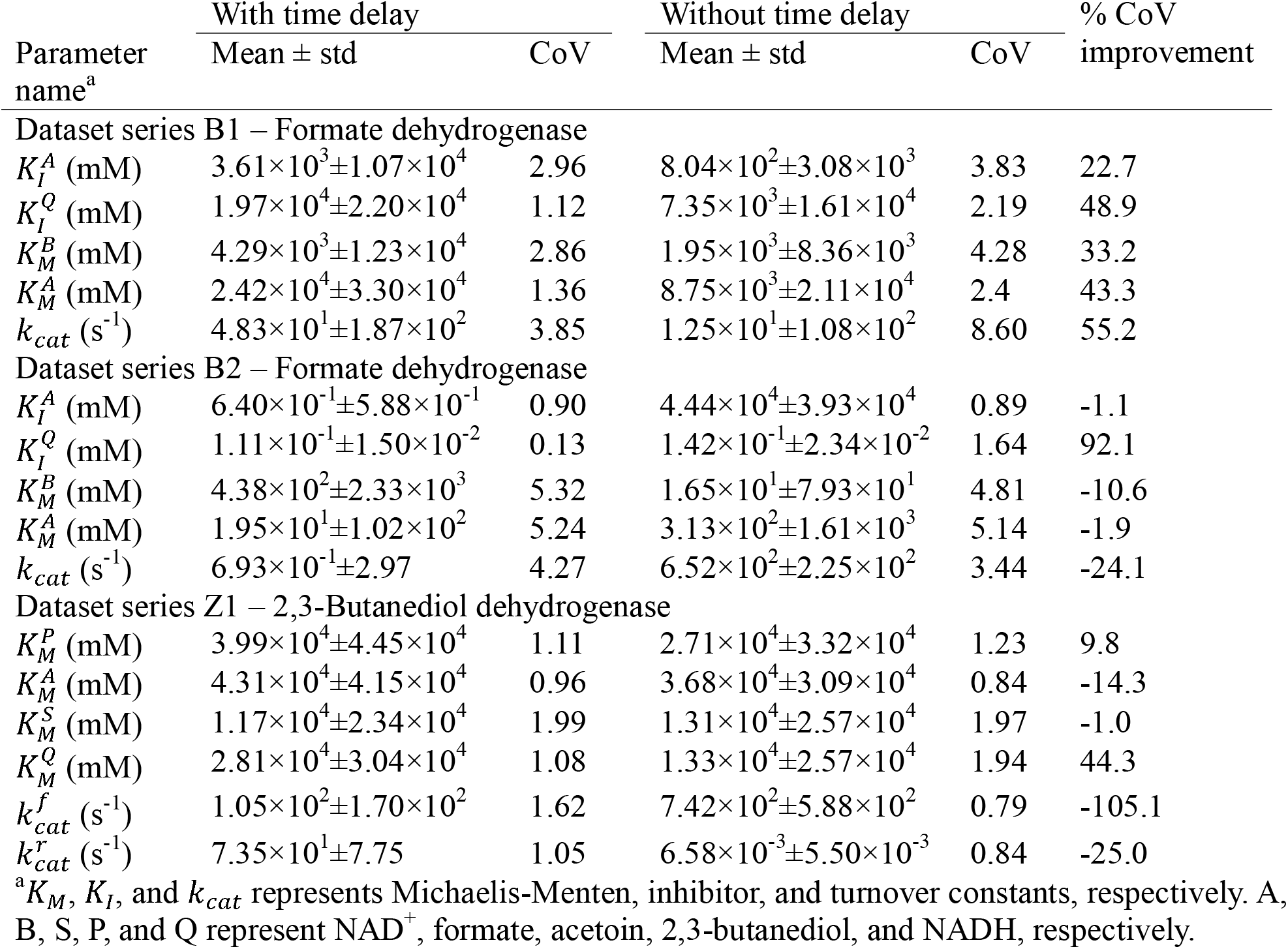
Mean, standard deviation (std), and coefficient of variation (CoV) of kinetic parameters found for dataset series B1, B2, and Z1.

Initial observation of predicted and experimental NADH time-course shows poor fitting where slight x-axis (time) shifts in experimental NADH concentrations occur (see Figure 3a). This shift is a systematic time-lag that was introduced into the dataset because there is an unaccounted time-lapse between the reaction start time (*i*.*e*., when the enzyme is mixed in with the metabolites) and the spectrophotometer measurement time. The resulting poor simulation fitting results from a mismatch of attempting to fit the datasets while fixing their initial measured concentrations. To correct this, KETCHUP’s utility function proactively searches via a bracketing search method to recommend an optimal time-lag that minimizes the SSR for each dataset while constraining the initial metabolite concentrations. Each dataset in series B1, B2, and Z1 are separately parameterized to find a corresponding time-lag parameter. Then each dataset series is re-parameterized with time-lag adjustments (see Supplementary Figures SF4-SF6). The implementation of time-lag adjustments was able to slightly improve SSR and solutions converged (See Table 2). However, time-lag parameters for each dataset in any given dataset series are distinct from one another indicating possible error in experimental setup resulting in uneven lags across datasets on top of possible measurement noise error.

**Figure 3.**
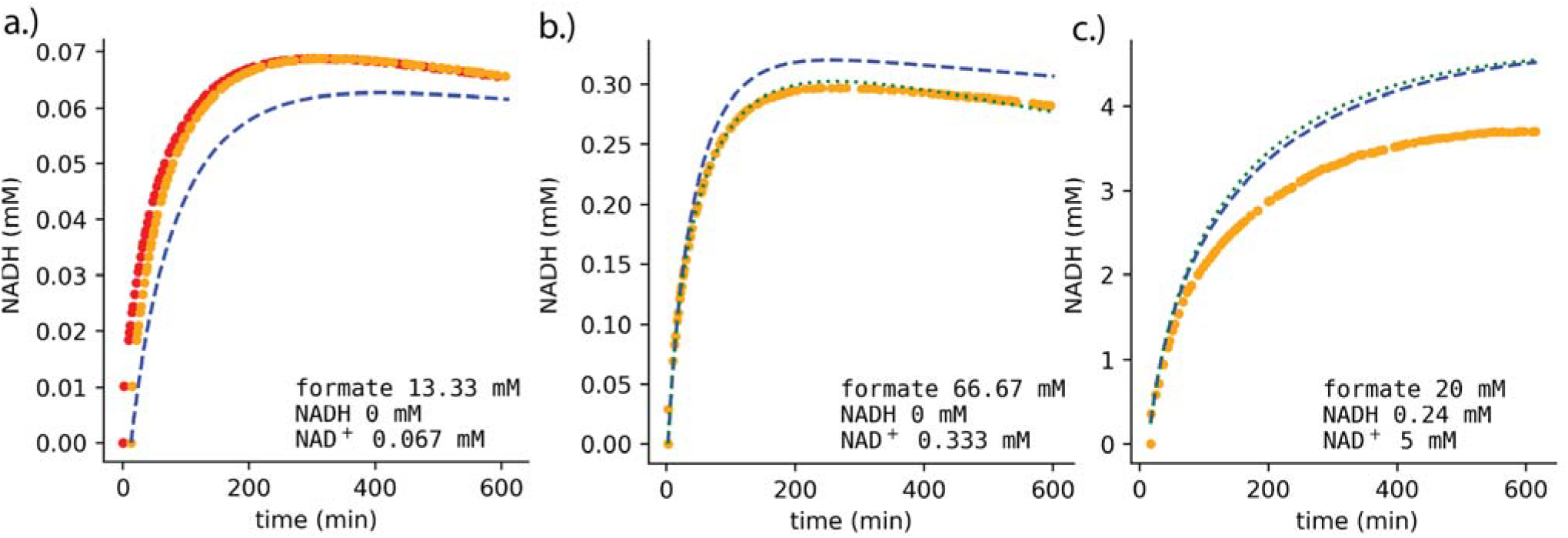
Prediction of NADH concentration for FDH over time between experimental (red dots), experimental with adjusted time delay (orange dots), simulation of separately parameterized dataset (dotted green line), and simulation of combined datasets (dashed blue line). Plots demonstrate a.) improvement of fit (SSR) with time-delay parameter introduced, b.) separately parameterized set verifying proposed mechanistic rate law can and c.) cannot fit dataset. Initial metabolite concentrations are in plot text. Plots of prediction and simulation of all datasets for both FDH and BDH in Supplementary Figures SF4-SF6.

**Figure 4.**
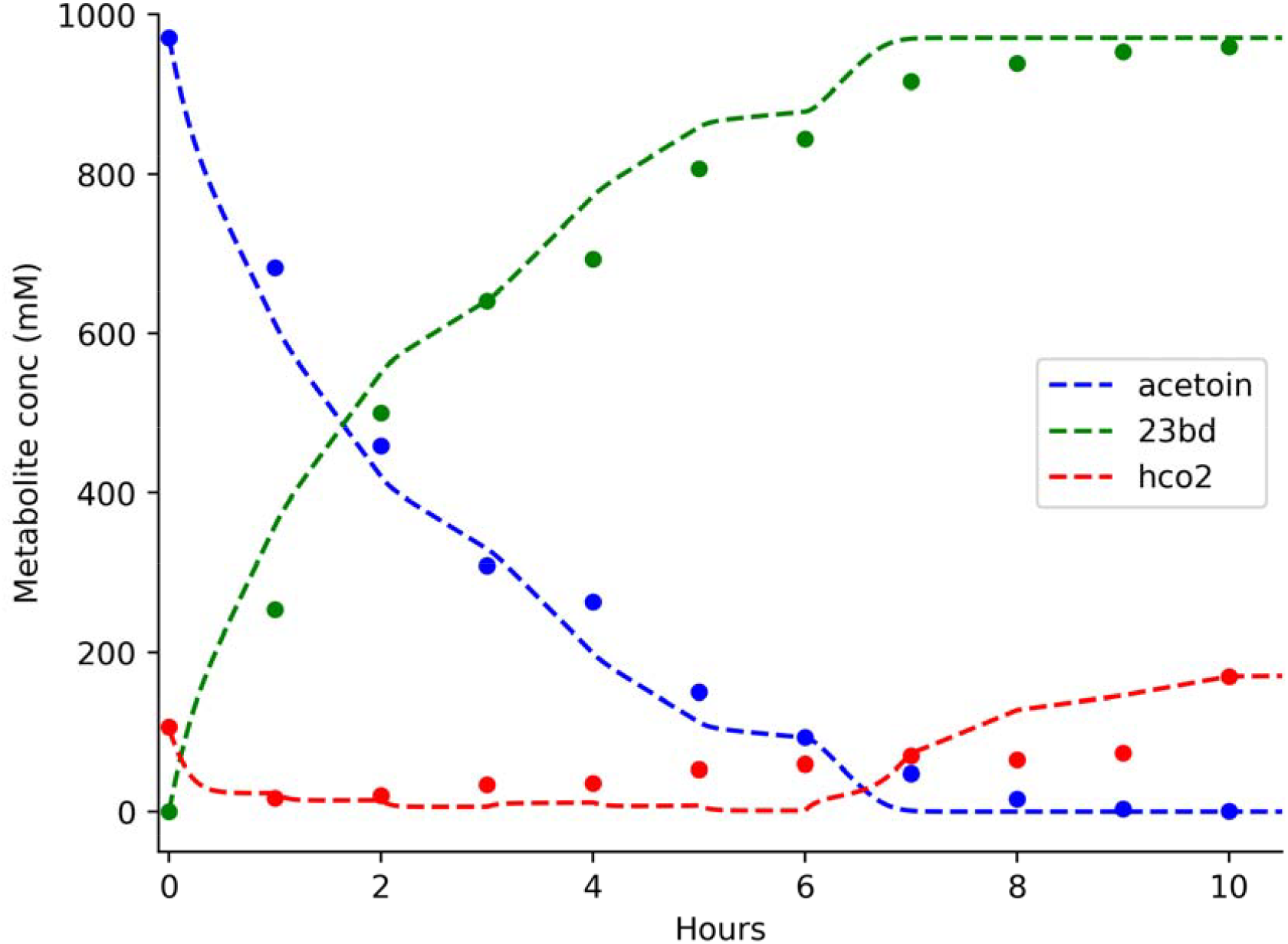
Prediction of Binary FDH-BDH system against experimental data (dotted points) and simulation (dashed lines). Simulation was performed using the best fitted solutions of single-enzyme parameterizations.

Time-lag adjustments also slightly improved kinetic parameter resolution (See Table 3). We determined kinetic parameter resolution by observing the coefficient of variation (CoV) which is the ratio of the standard deviation to the mean and used to identify the degree of variation in kinetic parameters [68]. There is substantial improvement for the only NADH involved kinetic parameter for both FDH (*i.e*., 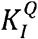 CoV of 1.12 vs 2.19 and 0.13 vs 1.64 for dataset series B1 and B2, respectively) and BDH (*i.e*., 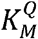 CoV of 1.08 vs 1.94), demonstrating the importance of time-delay adjustments for NADH profiles. This improvement highlights not only the importance of having well-aligned time-course data but also the inclusion of different metabolite data as well.

**Table 3.** Performance metrics for parameterization of dataset series B1, B2, and Z1.

| Dataset Series (Enzyme) | # of datasets in series (DF) | With time delay |  |  | Without time delay |  |  |
| --- | --- | --- | --- | --- | --- | --- | --- |
|  |  | Best SSR<sup>a</sup> (mM<sup>2</sup>) | % solutions converged | Average solve time (seconds) | Best SSR (mM<sup>2</sup>) | % solutions converged | Average solve time (seconds) |
| B1 (FDH) | 19 (7) | 1.94 | 95.6 | 137.55 | 2.31 | 81 | 148.80 |
| B2 (FDH) | 59 (7) | 162.70 | 63.6 | 674.40 | 194.61 | 16.4 | 766.62 |
| Z1 (BDH) | 78 (8) | 24.15 | 3.0 | 897.39 | 26.28 | 1.6 | 1378.40 |

<sup>a</sup>SSR value for each kinetic model solution falls within the $\chi^2$ statistic for a p-value of 0.05 with their corresponding degrees of freedom (DF).

After parameterization of kinetic models, we compare the fitting datasets to their simulation from *(a)* single-dataset and *(b)* full dataset series parameterized solution. A few simulations from scenario *(a)* show substantial deviation from the fitted datapoints (See example in Figure 3b). Normally, we would only expect that deviations occur in scenario (*b*) because the kinetic model is attempting to explain a variety of metabolite and enzyme concentrations with only one set of kinetic parameters. This discrepancy in scenario *(a)* likely results from either *(i)* poor selection of rate law representing the enzyme, *(ii)* lacking information on substrate or product inhibition, or (*iii)* noise in experimental measurements.

### Simulation of fed-batch binary system with single-enzyme kinetic parameters

To validate the assumption that kinetic parameters found in single-enzyme parameterizations can be used in multi-enzyme systems, an FDH-BDH binary system was simulated and compared against experimental results [69]. The initial experimental conditions are set to 971 mM of acetoin, 1 mM of NADH and 104 mM of formate measured with HPLC based on communications with the authors. Because the experiment implemented pH control (*i.e*., addition of formic acid) and removed volume for samples, experimental data used for comparison were re-normalized to millimolar units after accounting for volume change. Simulation of the binary system was performed in piecewise fashion with each time-step matching the duration (*i.e*., 60 minutes between sampling) between sampling timepoints with the previous end metabolite concentrations being reset as the initial condition in the next time-step. This piecewise simulation allows for flexible adjustment of formic acid addition rates to the simulation. Due to lack of pH control information, formate addition rate is assumed to be a linear between experimental datapoints (*e.g*., a formate addition rate of 4.63 mM/minutes was assumed between timepoints 0 and 60 minutes). Note that the varying formate addition rates result in “non-smooth” metabolite profiles. The simulation (using best solution kinetic parameters from both enzymes) was able to closely replicate the experimental metabolite profile (See Figure 4) and demonstrate ∼100% theoretical yield of 2,3-butanediol like the experimental results. Overall, this result demonstrates the feasibility of first resolving kinetic parameters for single enzymes and subsequently using the determined values to simulate multi-enzyme reaction cascades.

## Discussion

The parameterization of kinetic models for cell-free systems, specifically single-enzymes, offers potential in exploration and identification of rate laws and regulation to help better characterize each enzyme [48]. However, the parameterization of single-enzymes requires metabolite time-course data of varying initial concentrations. Herein, KETCHUP demonstrated fitting of single-enzyme datasets with time-lag correction and a resulting binary enzyme system, using single-enzyme fitted kinetic parameters, can closely simulate and replicate experimental metabolite profiles. These results demonstrate opportunities to efficiently characterize single enzymes in a high-throughput fashion for the systematic piecemeal construction of larger kinetic models that simulate cascade systems (*e.g*., combining four separate kinetic and enzyme information to create a quad-enzyme system converting pyruvate to 2,3-butanediol [69]).

In this study, we first benchmarked KETCHUP’s performance and kinetic parameter estimations with a previously studied enzyme and characterized mechanistic rate law, formate dehydrogenase from *C. boidinii*. The best parameterized model (*i.e*., model yielding the lowest SSR) was able to adequately recapitulate the fitted data. We observed a slight systematic overprediction in NADH time-course for lower initial NAD^+^ concentrations resulting from the objective of prioritizing kinetic parameters to better fit those with high NADH concentrations. This overprediction characteristic can be adjusted to consider both low and high concentration NADH profiles during fitting by including weights to normalize the objective function. However, it is important to carefully consider weights for each datapoint in a way that does not bias the overall fit (*e.g*., measurements for lower NADH concentrations may be within the range of experimental noise). An important advantage purified enzyme cell-free systems has over *vivo* systems is the straightforward reaction environment that allows for selective metabolite testing (*i.e*., addition/removal of known metabolites) for the observation of kinetic rates and allosteric regulations. Several factors can improve the resolution of kinetic parameters such as inclusion of different metabolite profiles and proper timed measurements. For example, NADH can be readily measured via spectrophotometer but CO_2_ (*i.e*., product of FDH) cannot be measured in 384 well-plates. For the latter factor, KETCHUP provides a method to correct time discrepancies in high-throughput measurements by actively searching for the most suitable time-lag parameter to best fit the datasets.

We parameterized FDH and BDH kinetic models capable of recapitulating most experimental datasets. Discrepancies between fits result from lacking metabolite profiles (except for NADH) and possibly proper rate laws used for modeling. For example, the published FDH rate law used did not include any Michaelis-Menten constants related to product metabolites (*i.e*., CO_2_ and NADH) assuming that CO_2_ is rapidly dispersed into gaseous phase which renders reaction FDH irreversible. However, this assumption is only true under very specific experimental conditions. For example, in the presence of active pH control (as we used in our two-enzyme FDH-BDH cascade reaction), the addition of concentrated acid causes rapid CO_2_ evolution from the liquid phase. On the other hand, when pH is controlled via buffering, the conversion of CO_2_ between its various carbonate forms is highly reversible and can and can affect FDH reversibility rates [70] and cascade reactions involving FDH [69].

For BDH, there is no published mechanistic rate law available so convenience kinetics [65] was used as a starting point because the rate law is capable of capturing enzyme saturation effects, reaction stoichiometries, and allosteric regulations while only requiring a small number of parameters. Discrepancies in the fitting of BDH data likely result from the experimental set-up. Because BDH consumes NADH and there is a temperature discrepancy during the initial stages of the experiment, NADH profiles display sigmoidal behavior where BDH activity gradually speeds up over time. This lag-phase hinders parameterization of BDH by obscuring the true initial NADH concentration for when enzyme assay reaches experimental temperature. By trimming the lag-phase out of the datasets and assuming no NADH consumption before the enzyme assay reaches temperature, KETCHUP was able to still successfully find kinetic parameters that can recapitulate most of the fitting datasets. Using the solutions from both FDH and BDH kinetic models, an FDH-BDH binary system kinetic model was developed, simulated, and evaluated against experimental data. This capability demonstrates the potential of hierarchically building kinetic models by first parameterizing one enzyme system at a time and subsequently integrating all parameterized enzyme kinetics into a single model to simulate the multi-reaction cascade.

Overall, cell-free systems are ideally positioned for rational and systematic enzyme characterizations for both rate laws and allostery exploration. Their high throughput combined with algorithmic methods to filter out noise allows for rapid parameterization of kinetic parameters. The extension introduced in this study serves as a pilot for parameterizing time-course data with KETCHUP, an open-source framework that was originally developed for scalable parameterization of large-scale kinetic systems using steady-state datasets with flexible objective function and customizable rate laws. This extension of KETCHUP enhances the design-test-build-learn cycle by demonstrating an automatable workflow that can speed-up decisions in experimental design (*e.g*., which metabolite profiles to capture or initial metabolite concentrations). These features allow for straightforward implementation of user-defined searches (*e.g*., finding the lowest set of kinetic parameters while minimizing SSR) and specific kinetic rate laws (*i.e*., user-defined FDH reaction in this study). This tool’s adaptability towards user selections facilitates the semi-automatic construction and parameterization of customizable kinetic models compared to existing options such as MATLAB, which requires licensing or COPASI defaults kinetic rate-law selections to either mass action kinetics or simple Michaelis-Menten equations. Development of these kinetic models to characterize enzyme using cell-free systems serves as a platform for the development of detailed large-scale kinetic models and application of these models towards engineering *in vivo* systems.

## Materials and methods

### Data collection

In this study, we collected NADH time-course data for *C. boidinii* FDH enzyme purchased from Sigma (Roche) (*i.e*., Dataset series B1 and B2) and purified BDH protein sequenced from *S. marcescen* (*i.e*., Dataset series Z1). Dataset series vary by initial substrate and enzyme concentrations ranges.

Individual enzyme activity measurements were performed in a 384 well plate in BioTek plate reader at 37°C. Activity was determined based on changes in the concentration of NADH, determined spectrophotometrically at 340 nm. In some cases, additional measurements at 390 nm and 400 nm were used to extend the dynamic range of measurements. NADH concentrations (in mM) are provided in Supplementary Materials SM1 for Datasets series B1, B2, and Z1. The absorption values were converted to NADH concentration using the following extinction coefficients: 6.220 mM/Abs_340_/cm, 0.4276 mM/Abs_390_/cm, 0.1205 mM/Abs_400_/cm. The Abs_340_ extinction coefficient is a well-established constant [71]. The values at 390 and 400 nm were determined empirically. For a 60 μl reaction in a 384 well plate, the pathlength was 0.623 cm.

### Purification of BDH protein

Purification of the BDH protein has been described in detail previously [69]. Briefly, *E. coli* BL21 DE3 harboring 6x tagged was cultured overnight in LB medium in the presence of ampicillin (100 μg/mL). The secondary culture was started using 1% of the saturated overnight culture and induced using 0.3 mM IPTG at OD_600_ of ∼ 0.4 after cultivation at 37 °C. Post induction, cells were harvested after 16 h cultivation at 18 °C and stored in -80 °C overnight. Pellets were resuspended in lysis buffer consisting of 20 mM tris-HCl pH 7.5, 500 mM NaCl, 5 mM imidazole, 1 mg/mL lysozyme and 1mM phenylmethylsulfonyl fluoride. Cells were lysed using microtip sonicator. After centrifugation of lysate at 9,000*g* (1 h and 4 °C), the supernatant was filtered (0.45 μm) and purified via Ni-nitrilotriacetic acid (Ni-NTA) metal affinity chromatography. The components of wash buffer were 20 mM tris-HCl pH 7.5, 500 mM NaCl and 15 mM imidazole. Protein elution was performed by buffer consisting of 20 mM tris-HCl pH 7.5, 500 mM NaCl and 650 mM imidazole. The purified protein was dialyzed in 100 mM sodium phosphate buffer and purity checked on SDS-PAGE gel. The concentration of BDH was determined by bicinchoninic acid (BCA) protein assay kit (G-Biosciences, MO, USA).

### Time-lag adjustments

Dynamic data often needs to be corrected for a time-lag that occurs due to the difference between when the assay is started (by addition of the starting reagent) and when the data collection starts. This time-lag represents the amount of time required to add and mix the starting reagent, apply the sealing film (if used to prevent evaporation), load the plate into the plate reader instrument, and initiate the data collection. Typical values range from 10 to 60 seconds.

To identify the correct time lag (in seconds) for each dataset, a bracketing search method is used to iteratively search for an optimal time lag time parameter that minimizes the SSR of each dataset (See below). After the level of all time lags have been identified for each dataset, the value of each time lag parameter is fixed for all subsequent simulations. A kinetic model is parameterized using the time lags to improve fit and resolve parameters estimated.

An initial guess of time lag (*t*^0^) is first predicted by extrapolating the measured NADH concentration back to its initial value by fitting the data points to a cubic polynomial and then identifying the root value closest to zero (for FDH) or [NADH_init_] for BDH. This fitting is done using the NumPy package [72] and functions polyfit and roots.

STEP 1: Initialize counter k = 0, time lag search interval step size *i*^0^ = *t*^0^, stop criteria *ϵ*= 0.001.

STEP 2: Set time lag search interval [*t*^*k*^ − *i*^*k*^, *t*^*k*^ + *i*^*k*^].

STEP 3: Divide search interval into ten evenly spaced search times G with step size *s*^0^ such that,

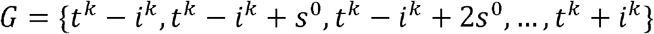

STEP 4: For *g ϵ G*, run KETCHUP with 20 multi-starts and record lowest SSR 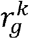 and corresponding time lag 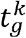.

STEP 5: If lowest SSR lower than best SSR, update best SSR *r*^*best*^ and corresponding time lag value *t*^*best*^.

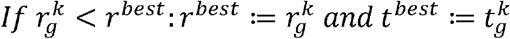

STEP 6: Update time lag search interval step size.

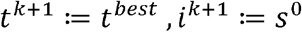

STEP 7: Check time interval step size.

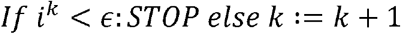

### Model parameterization using time-course data

Because KETCHUP was originally developed to fit steady-state data, the Pyomo differential algebraic equation (DAE) package [73] was used to discretize the rate laws which are in the form of ordinary differential equations (ODE). This is done by first declaring metabolite concentrations (*i.e*., NADH) as a derivative variable (*i.e*., Pyomo DAE DerivativeVar component) and time as a continuous set (*i.e*., Pyomo DAE ContinuousSet component). Once the rate law is set to their respective ODE, Pyomo DAE discretizes the ODE using the Implicit Euler method (*i.e*., default setting of Pyomo DAE discretizer) with the number of finite elements equal the number of time-points for the respective dataset. Pyomo is then used to formulate the remaining stoichiometric and feasibility (*e.g*., non-negative kinetic parameters and concentrations) constraints. The objective function is set to minimize the SSR between predicted and fitted data with an equal weighting of one for each data point (See Equation 1). Parameterization is then carried out with the nonlinear programming solver, Interior Point Optimizer (IPOPT [74]) with default threshold settings (See [62] for full details on IPOPT implementation in KETCHUP). Simulation of metabolite concentrations, for single and binary enzymes, are implemented using Pyomo DAE’s built-in simulator package which utilizes the CasADi framework [75] which uses the Implicit Differential-Algebraic Solver (IDA) [76] to solve the differential-algebraic equations defined by the rate laws.

*Equation 1: Sum of squared residual (SSR) for minimizing objective function used in the parameterization of kinetic models with time-course data, where t is time point, y is the measured experimental (meas) and model predicted (pred) values, and k is the index of dataset used during parameterization for all datasets K*.

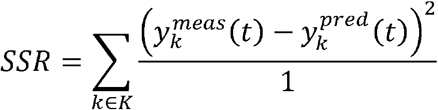

### Computational implementation

KETCHUP is programmed in Python v3.8.5 and makes use of library packages provided through Anaconda. Formulation of equations and constraints are done through Pyomo v6.4.0. Parameterization and solving of equations are done with IPOPT v3.14.0, which interfaces with Pyomo, was recompiled from source code using GNU compiler v8.3.1 [77] to include linear solver ma97 [78] from the Harwell Subroutine Library [79]. Computations were performed on dual 10-or 12-core Xeon E5-2680 processors with InfiniBand using 1 core and 4 GB RAM running Red Hat Enterprise Linux Server release 7.9. KETCHUP uses COBRApy v0.25.0 [80] for reaction and metabolite parsing, Pandas v1.2.25 [81] for internal data structures and storing experimental data, and NumPy v1.23.1 [72] to randomize initial kinetic parameters. Converged kinetic model solutions can be exported to SBML [82] format using the libSBML package [83]. A generated graphical user interface is developed using Streamlit [84].

## Conflict-of-interest statement

The authors declare no commercial or financial conflict of interest.

### CRediT authorship contribution statement

**Mengqi Hu:** Writing – review & editing, Writing – original draft, visualization, validation, Software, Methodology, Formal analysis, Data curation

**Bilal S. Jilani:** Data curation, Formal analysis, Validation, Investigation, Writing – original draft **Daniel G. Olson:** Funding acquisition, Project administration, Conceptualization, Supervision, Resources, Investigation, Data curation, Formal analysis, Writing – review & editing

**Costas D. Maranas:** Funding acquisition, Conceptualization, Supervision, Resources, Writing – review & editing

## Acknowledgements

The authors wish to acknowledge Dr. Wheaton L. Schroeder at Washington State University for useful discussions on data processing and Dr. Patrick F. Suthers at The Pennsylvania State University for informative discussion on the implementation of time-lag algorithm and in coding of KETCHUP. Computations for this research were performed on the Pennsylvania State University’s Institute for Computational and Data Sciences’ RoarCollab supercomputer.

