## Supplementary Figures for "Parameterization of cell-free systems with time-series data using KETCHUP": Supplementary Figures.docx

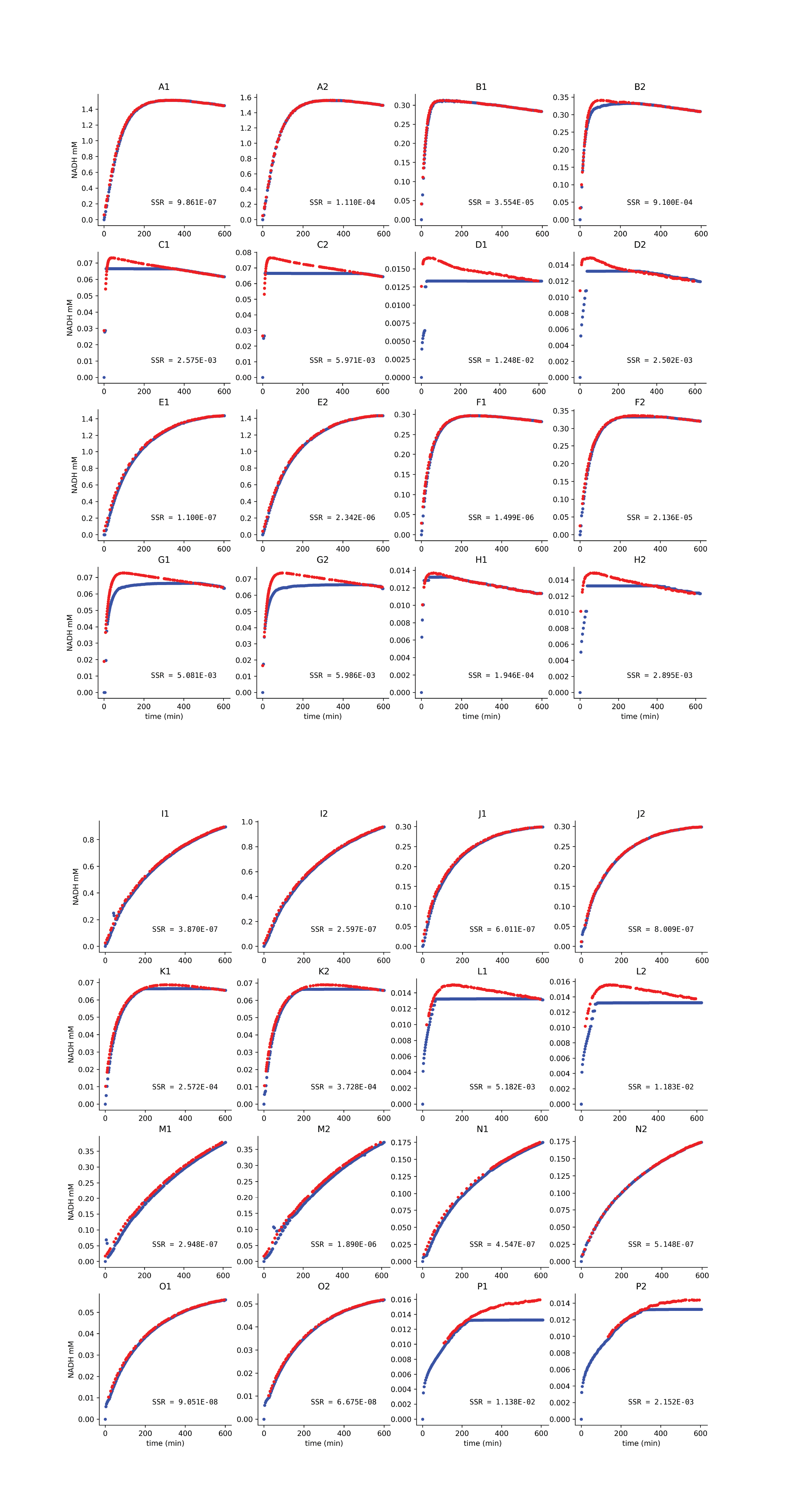

*Supplementary Figure SF1: Raw data (red dots) and predicted (blue dots) for Formate dehydrogenase dataset B1. Each plot is labeled with their respective well id and SSR fit value from parameterization of the single dataset.*

*Supplementary Figure SF2: Raw data (red dots) and predicted (blue dots) for Formate dehydrogenase dataset B2. Each plot is labeled with their respective well id and SSR fit value from parameterization of the single dataset.*

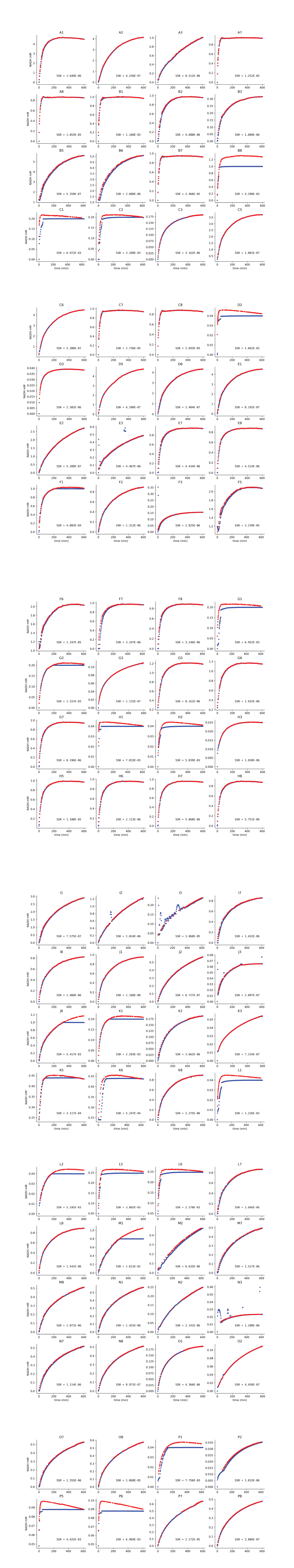

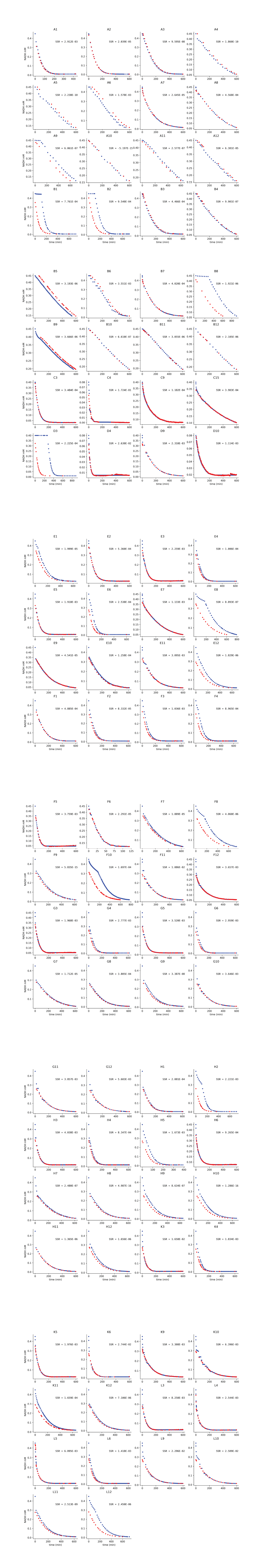

*Supplementary Figure SF3: Raw data (red dots) and predicted (blue dots) for 2,3-butanediol dehydrogenase dataset Z1. Each plot is labeled with their respective well id and SSR fit value from parameterization of the single dataset.*

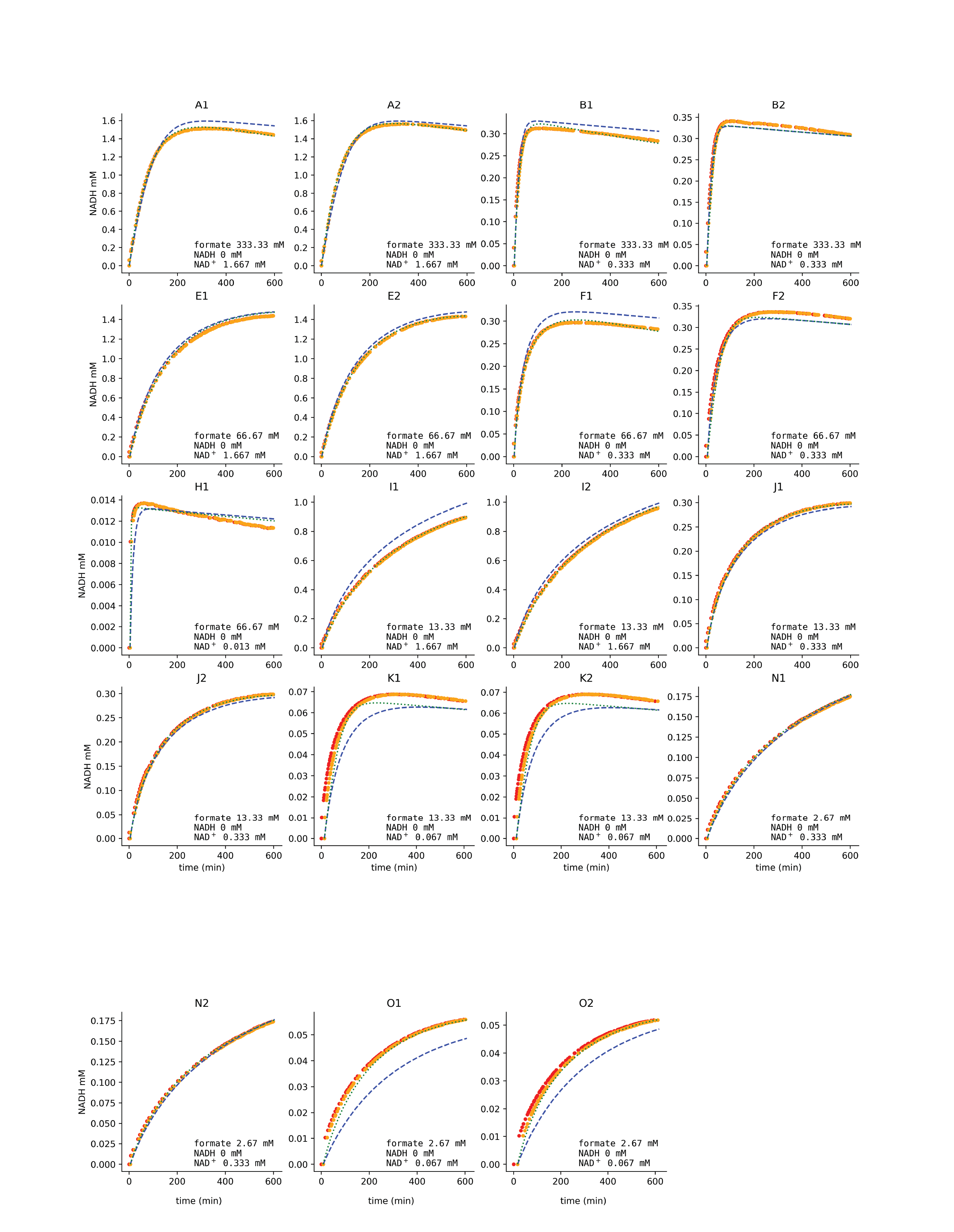
*Supplementary Figure SF4: Simulation of NADH concentration for FDH dataset B1 over time between experimental (red dots), experimental with adjusted time delay (orange dots), simulation of separately parameterized dataset (dotted green line), and simulation of combined datasets (dashed blue line). Plot contains text of initial metabolite information and respective experimental well as title.*

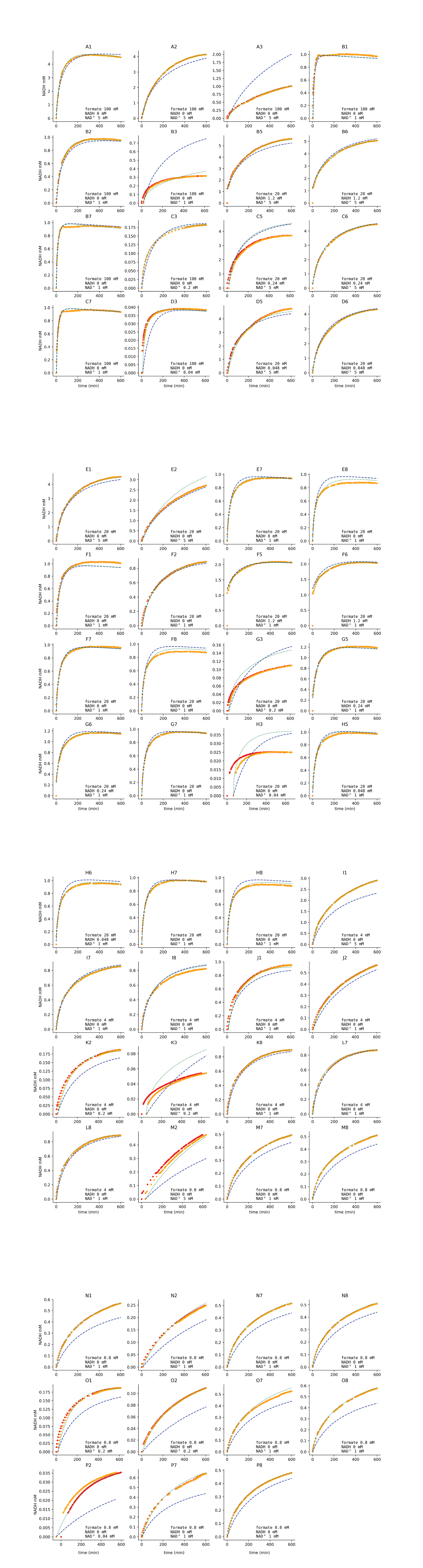
*Supplementary Figure SF5: Simulation of NADH concentration for FDH dataset B2 over time between experimental (red dots), experimental with adjusted time delay (orange dots), simulation of separately parameterized dataset (dotted green line), and simulation of combined datasets (dashed blue line). Plot contains text of initial metabolite information and respective experimental well as title.*

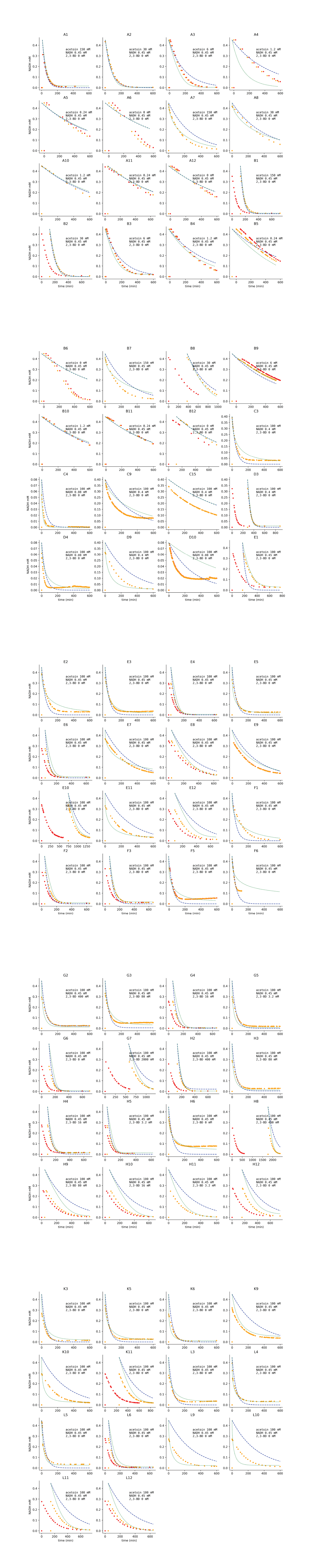
 *Supplementary Figure SF6: Simulation of NADH concentration for BDH dataset Z1 over time between experimental (red dots), experimental with adjusted time delay (orange dots), simulation of separately parameterized dataset (dotted green line), and simulation of combined datasets (dashed blue line). Plot contains text of initial metabolite information and respective experimental well as title.*
