## Supplementary Materials SM2 for "Parameterization of cell-free systems with time-series data using KETCHUP": Data Thinning (BDH example).ipynb

```
# %% [code]
import pandas as pd
import matplotlib.pyplot as plt
import numpy as np
from rdp import rdp
from matplotlib.offsetbox import AnchoredText

# %% [code]
xl = pd.ExcelFile('.\data\BDH_data_from_Olson-2022-06.xlsx')
sheetnames = xl.sheet_names

# %% [code]
c = 0; already_plotted = []
ma_window = 10
writer =  pd.ExcelWriter("BDH_data_from_Bilal_2022-11_filtered.xlsx")
for s in ["A2"]:#sheetnames:
    if c % 9== 0:
        fig, ((ax1,ax2,ax3),(ax4,ax5,ax6),(ax7,ax8,ax9)) = plt.subplots(3,3,figsize = (15,15))
        ax = [ax1,ax2,ax3,ax4,ax5,ax6,ax7,ax8,ax9]

    df = pd.read_excel('.\data\BDH_data_from_Olson-2022-06.xlsx', skiprows=[0,1,2,3,4,5,6],sheet_name = s)
    df2 = pd.read_excel('.\data\BDH_data_from_Olson-2022-06.xlsx', sheet_name = s,header=None)
    first_col = df2.loc[0:7,0].tolist()
    second_col = df2.loc[0:7,1].tolist() 
    time_title = 'Time (mins)'
    nadh_conc = second_col[5]
    index = 0
    for i,j in enumerate(df['NADH'].tolist()):
        if j > nadh_conc: index = i + 1
    df = df.iloc[index:,:]

    try:
        ax[c%9].plot(df[time_title],df['NADH'])
    except KeyError:
        time_title = 'time'
        ax[c%9].plot(df[time_title],df['NADH'])

    #take the moving average
    move_av = df.rolling(window=ma_window).mean().dropna()
    move_av.reset_index(drop=True, inplace=True)

    #use the moving average instead to clean the data a little more
    arr = [];
    for i,d in enumerate(move_av[time_title].tolist()):
        arr.append(move_av.loc[i].tolist())
    arr.insert(0,df.iloc[0].tolist()) #retains first row data that is removed when using rolling window
        
    target_data_pts = 100
    multiplier = 1
    for i in range(500):
        den = 1000
        den *= multiplier
        eps = (max(move_av['NADH'].tolist()) - min(move_av['NADH'].tolist()))/den
        filtered_data = rdp(np.array(arr), algo='iter',epsilon=eps)
        datapts = len(filtered_data)
        if datapts <= target_data_pts + 5 and datapts >= target_data_pts - 5:
            break
        if datapts < target_data_pts: multiplier += 0.10
        if datapts > target_data_pts: multiplier -= 0.01
            
    ax[c%9].plot(filtered_data[:,0],filtered_data[:,1], color='red')
    
    ##root finding - 3rd degree
    #poly fit
    x = filtered_data[:,0]
    y = filtered_data[:,1]
    
    #filter data by datapoints
    x2 = []; y2 = []; diff = max(y) - min(y);threshold = diff * 0.05
    for i,_ in enumerate(x):
        if i != 0 and abs(y[i] - y[i-1]) < threshold: break
        x2.append(x[i]);y2.append(y[i])
    #filter data by number of data points    
    x3 = x[:1*len(x)//3];y3 = y[:1*len(y)//3]

    xfit = x2 if len(x2) > len(x3) else x3
    yfit = y2 if len(y2) > len(y3) else y3
    
    trend = np.polyfit(xfit,yfit,3)
    trend[3] = trend[3] - nadh_conc
    trendpoly = np.poly1d(trend)
    #find root
    r = np.roots(trend)
    root = round(min(r.real[abs(r.imag)<1e-5], key = abs),3)
    ax[c%9].plot(xfit,trendpoly(xfit)+nadh_conc, color = 'green')
    text = f'estimated root value : {root}'
    ax[c%9].add_artist(AnchoredText(text,loc='upper right'))
    
    for d in filtered_data[:,0]:
        first_col.append(d)
    for d in filtered_data[:,1]:
        second_col.append(d)

    #write epsilon in scientific notation
    eps_sci = ("{:.3e}".format(eps))

    ax[c%9].set_title(f"{s} orig: {len(move_av['NADH'].tolist())} | filt: {len(filtered_data)} data pts \n {round((1 - len(filtered_data)/len(move_av['NADH'].tolist()))*100,2)}% removed | epsilon = {eps_sci}")
    c += 1
    
    if c % 9 == 0: plt.show()
    
    pd.DataFrame([first_col,second_col]).transpose().to_excel(writer,sheet_name=s, index=False,header=False)

writer.close()
xl.close()
writer.handles=None
plt.show()
```
