## Supplementary Materials SM2 for "Parameterization of cell-free systems with time-series data using KETCHUP": simulation binary system.ipynb

```
# %% [code]
import os, sys, json
from pyomo.environ import *
from pyomo.dae import *
from pyomo.dae.simulator import Simulator
import numpy as np
import logging
logging.getLogger('pyomo.core').setLevel(logging.ERROR)
import matplotlib.pyplot as plt
exp_data = {
    'time':[0,1,2,3,4,5,6,7,8,9,10,24],
    'acetoin':[971.0439913,682.6181848,459.208027,308.3003315,263.2544518,149.8680396,
               93.46151024,47.84479588,16.04114794,3.147777154,0.5864201,0],
    '23bd':[0,253.390451,499.7211577,640.7983832,692.9552927,806.5389984,843.7616898,
                915.953227,938.4841003,953.1739333,959.6760942,975.8266099],
    'hco2':[106.4470588,17.11434343,20.22083859,33.91695502,35.67344451,52.71546131,
                59.38795452,69.70842223,64.98969826,73.3982515,169.3867239,188.6024144]
}

# %% [code]
def create_model(maxtime,form_add_rate):
    with open('BDH_optimal_results_data_153.txt') as f:
        bdh_res = json.load(f)
    with open('FDH_optimal_results_data_272.txt') as f:
        fdh_res = json.load(f)

    model = ConcreteModel()
    #BDH variables
    del bdh_res['kp']['kcat_f']['NADH_decomp']
    del bdh_res['kp']['kcat_r']['NADH_decomp']
    model.KM_products = Param(Set(initialize=['BDH+23bd','BDH+nad']),initialize=bdh_res['kp']['Km_p'])
    model.KM_reactants = Param(Set(initialize=['BDH+acetoin','BDH+nadh','NADH_decomp+nadh']),initialize=bdh_res['kp']['Km_r'])
    model.Kcat_f = Param(Set(initialize=['BDH']),initialize=bdh_res['kp']['kcat_f'])
    model.Kcat_r = Param(Set(initialize=['BDH']),initialize=bdh_res['kp']['kcat_r'])
    #FDH initialization
    model.kcat = Param(initialize=fdh_res['kp']['kcat']['null'])
    model.Km_r = Param(Set(initialize = ["nad","hco2"]), initialize=fdh_res['kp']['Km_r'])
    model.K_decomp = Param(initialize=bdh_res['kp']['K_decomp']['null'])
    model.Ki = Param(Set(initialize=['nad','nadh']),initialize=fdh_res['kp']['Ki'])
    #define components
    model.TIME = ContinuousSet(bounds=(0,maxtime))
    model.c = Var(model.TIME, Set(initialize = ['acetoin', 'nadh', 'nad', '23bd', 'hco2', 'co2']))
    model.fdh_rate = Var(model.TIME)
    model.bdh_rate = Var(model.TIME)
    model.nadh_decomp = Var(model.TIME)
    model.dcdt = DerivativeVar(model.c, wrt=model.TIME, within=Reals)
    model.e = Var(model.TIME, Set(initialize=['BDH','FDH']))
    #dcdt
    def dcdt_acetoin(m,t):
        return m.dcdt[t,'acetoin'] == - m.bdh_rate[t]
    model.diff_acetoin = Constraint(model.TIME,rule=dcdt_acetoin)
    def dcdt_nadh(m,t):
        return m.dcdt[t,'nadh'] == -m.bdh_rate[t]  - m.nadh_decomp[t] + m.fdh_rate[t]
    model.diff_nadh = Constraint(model.TIME,rule=dcdt_nadh)
    def dcdt_nad(m,t):
        return m.dcdt[t,'nad'] == m.bdh_rate[t] - m.fdh_rate[t]
    model.diff_nad = Constraint(model.TIME,rule=dcdt_nad)
    def dcdt_23bd(m,t):
        return m.dcdt[t,'23bd'] == m.bdh_rate[t]
    model.diff_23bd = Constraint(model.TIME,rule=dcdt_23bd)
    def dcdt_hco2(m,t):
        return m.dcdt[t,'hco2'] ==  -m.fdh_rate[t] + form_add_rate
    model.diff_hco2 = Constraint(model.TIME,rule=dcdt_hco2)
    def dcdt_co2(m,t):
        return m.dcdt[t,'co2'] == m.fdh_rate[t]
    model.diff_co2 = Constraint(model.TIME,rule=dcdt_co2)
    #define rates
    def bdh_rate(m,t):
        num = (m.Kcat_f['BDH']*(m.c[t,'acetoin']*m.c[t,'nadh'])/(m.KM_reactants['BDH+acetoin']*m.KM_reactants['BDH+nadh'])- 
               m.Kcat_r['BDH']*(m.c[t,'23bd']*m.c[t,'nad'])/(m.KM_products['BDH+23bd']*m.KM_products['BDH+nad']))
        den = (1+m.c[t,'acetoin']/m.KM_reactants['BDH+acetoin'])*(1+ m.c[t,'nadh']/m.KM_reactants['BDH+nadh']) + \
               (1+ m.c[t,'23bd']/m.KM_products['BDH+23bd'])*(1+ m.c[t,'nad']/m.KM_products['BDH+nad'])
        return m.bdh_rate[t] == num/den * m.e[t,'BDH']        
    model.bdh_rxn_rate = Constraint(model.TIME, rule=bdh_rate)
    def fdh_rate(m,t):
        num = m.kcat * m.c[t,'nad'] * m.c[t,'hco2'] * m.e[t,'FDH']
        den = m.Ki['nad'] * m.Km_r['hco2'] + \
              m.Km_r['hco2'] * m.c[t,'nad'] + \
              m.Km_r['nad'] * m.c[t,'hco2'] + \
              m.c[t,'nad'] * m.c[t,'hco2'] + \
              (m.Ki['nad']*m.Km_r['hco2'])/(m.Ki['nadh'])*m.c[t,'nadh'] + \
              (m.Km_r['nad']/m.Ki['nadh']) * m.c[t,'hco2'] * m.c[t,'nadh']
        return m.fdh_rate[t] == num/den 
    model.fdh_rxn_rate = Constraint(model.TIME, rule=fdh_rate)
    def nadh_decomp(m,t):
        return m.nadh_decomp[t] == m.K_decomp * m.c[t,'nadh']
    model.nadh_decomp_rate = Constraint(model.TIME, rule=nadh_decomp)
    def FDH_conc(m,t):
        #return m.e[t,'FDH'] == 0.0198 / (1.0899e-6)
        return m.e[t,'FDH'] == 0.0198 / (2.51e-3)
    model.fdh_add = Constraint(model.TIME, rule=FDH_conc)
    def BDH_conc(m,t):
        return m.e[t,'BDH'] == 0.0069 / (1.4131e-7)
    model.bdh_add = Constraint(model.TIME, rule=BDH_conc)

    return model

# %% [code]
def simulate_model(m):
    # Simulate the model using casadi
    sim = Simulator(m, package='casadi')
    tsim, profiles = sim.simulate(numpoints=60, integrator='idas')

    # Discretize model using Orthogonal Collocation
    discretizer = TransformationFactory('dae.finite_difference')
    discretizer.apply_to(model,wrt=model.TIME,nfe=60,scheme='BACKWARD')

    # Initialize the discretized model using the simulator profiles
    sim.initialize_model()

    return sim, tsim, profiles

def plot_result(m, sim, tsim, profiles):
    import matplotlib.pyplot as plt

    time = list(m.TIME)

    varorder = sim.get_variable_order()
    algorder = sim.get_variable_order(vartype='algebraic')

    for idx1, v in enumerate(varorder):
        plt.plot(tsim, profiles[:, idx1], label=v)

   # for idx2, v in enumerate(algorder):
   #     plt.plot(tsim, profiles[:, len(varorder) + idx2], label=v)


    plt.xlabel('t')
    plt.legend(loc='best')
    plt.show()

# %% [code]
time = []
concentrations = {'acetoin':[],'nadh':[],'nad':[],'23bd':[],'hco2':[],'co2':[]}
form_add_rates = {
    60:4.628301761,
    120:3.047268338,
    180:1.373303847,
    240:2.26983505,
    300:1.377732211,
    360:0.218306048,
    420:2.746208048,
    480:0.920658599,
    540:0.301448878,
    600:0.398002298,
    1440:0.02266383
}

for_total = [103.786]
lasttime = 0
for t in list(form_add_rates):
    
    #initial conditions
    if t == 60:
        model = create_model(60,form_add_rates[60])
        model.c[0,'acetoin'] = 971.04399
        model.c[0,'nadh'] = 1
        model.c[0,'nad'] = 0
        model.c[0,'23bd'] = 0
        model.c[0,'hco2'] = 103.786
        model.c[0,'co2'] = 0
    else:   
        model = create_model(t-lasttime,form_add_rates[t])
        #initial conditions carryover
        model.c[0,'acetoin'] = profiles[:,0][-1]
        model.c[0,'nadh'] = profiles[:,1][-1]
        model.c[0,'nad'] = profiles[:,2][-1]
        model.c[0,'23bd'] = profiles[:,3][-1]
        model.c[0,'hco2'] = profiles[:,4][-1]
        model.c[0,'co2'] = profiles[:,5][-1]
        
    sim, tsim, profiles = simulate_model(model)

    
    time += [t + lasttime for t in list(tsim)]    
    lasttime = 60 if t == 60 else t
    concentrations['acetoin'] = concentrations['acetoin'] + list(profiles[:,0])
    concentrations['nadh'] = concentrations['nadh'] + list(profiles[:,1])
    concentrations['nad'] = concentrations['nad'] + list(profiles[:,2])
    concentrations['23bd'] = concentrations['23bd'] + list(profiles[:,3])
    concentrations['hco2'] = concentrations['hco2'] + list(profiles[:,4])
    concentrations['co2'] = concentrations['co2'] + list(profiles[:,5])
    for_total.append(for_total[-1] + list(profiles[:,4])[-1])


# %% [code]
import matplotlib.pyplot as plt
colors = {'acetoin':'blue',
          'hco2':'red',
          '23bd':'green',
          'nadh':'orange',
          'nad':'purple',
          'co2':'yellow'
         }
for i,met in enumerate(concentrations):
    if met in ['acetoin','hco2','23bd']:
        plt.plot([t/60 for t in time], concentrations[met], label=met, color=colors[met],linestyle='--')
plt.scatter(exp_data['time'],exp_data['acetoin'],color=colors['acetoin'],s=20)
plt.scatter(exp_data['time'],exp_data['23bd'],color=colors['23bd'],s=20)
plt.scatter(exp_data['time'],exp_data['hco2'],color=colors['hco2'],s=20)
#plt.plot(exp_data['time'],for_total,color='orange',linestyle='--')
plt.xlabel('Hours')
plt.ylabel('Metabolite conc (mM)')
plt.xlim(0,12)
plt.ylim(-50,1000)
plt.legend(loc='best')
plt.show()

# %% [code]
with open('FDH_optimal_results_data_272.txt') as f:
    res1 = json.load(f)
with open('FDH_optimal_results_data_437.txt') as f:
    res2 = json.load(f)

# %% [code]
res1['kp']

# %% [code]
res2['kp']

# %% [code]
```
